# Genome-wide disruption of DNA methylation by 5-aza-2’-deoxycytidine in a parasitoid wasp

**DOI:** 10.1101/437202

**Authors:** Nicola Cook, Darren J Parker, Frances Turner, Eran Tauber, Bart A Pannebakker, David M Shuker

**Author notes:** These authors contributed equally (Nicola Cook, Darren J Parker).

## Abstract

DNA methylation of cytosine residues across the genome influences how many genes and phenotypes are regulated. As such, understanding the role of DNA methylation and other epigenetic mechanisms has become very much a part of mapping genotype to phenotype, a major question in evolutionary biology. Ideally, we would like to manipulate DNA methylation patterns on a genome-wide scale, to elucidate the role of epigenetic modifications in phenotypic expression. Recently, the demethylating agent 5-aza-2’-deoxycytidine (5-aza-dC; commonly used in the epigenetic treatment of certain cancers), has been deployed to explore the epigenetic regulation of a number of traits of interest to evolutionary ecologists. Recently, we showed that treatment with 5-aza-dC shifted patterns of sex allocation as predicted by genomic conflict theory in the parasitoid wasp *Nasonia vitripennis*. This was the first (albeit indirect) experimental evidence for genomic conflict over sex allocation facilitated by DNA methylation. However, this work lacked confirmation of the effects of 5-aza-dC on DNA methylation, drawing commentary on the efficacy of 5-aza-dC in a novel system. Here, using whole-genome bisulphite sequencing, we demonstrate unequivocally that 5-aza-dC disrupts methylation across the *Nasonia vitripennis* genome. We show that disruption leads to both hypo- and hyper-methylation, may vary across tissues and time of sampling, and that the effects of 5-aza-dC are context- and sequence specific. We conclude that 5-aza-dC has the potential to be repurposed as a tool in evolutionary ecology for studying the role of DNA methylation.

## Introduction

The genomics revolution has transformed our understanding of how genomes build phenotypes. The extraordinary diversity of morphologies, behaviours, and physiological adaptations we see across the natural world are not necessarily the product of different sets of genes, or different numbers of genes in organisms, but rather differences in how, where and when genes are expressed during development. The genomics revolution has therefore been as much about understanding gene regulation as it has been about identifying allelic variants at genetic loci (Biémont 2010, Ashbrook *et al.* 2018). At the forefront of our growing knowledge of gene regulation has been the role of epigenetic modification of DNA, including DNA methylation. In the vast majority of cases, DNA methylation occurs at the C5 position of the cytosine ring in CpG dinucleotides and is catalysed by DNA methyltransferase enzymes (Dnmts). Three types of Dnmt are known in eukaryotes; Dnmt3 establishes DNA methylation patterns *de novo*, Dnmt1 maintains these patterns, and Dnmt2 is involved in tRNA methylation (Lyko 2018). In mammals, broadly, DNA methylation is found throughout the genome (except at CpG islands near the promoters of genes) and is associated with transcriptional repression (Feng *et al.* 2010, Zemach *et al.* 2010). By contrast, in insects, DNA methylation is concentrated in gene bodies and is associated with more stable patterns of gene expression (Bewick *et al.* 2017). Notably, vertebrate and invertebrate DNA methylation varies greatly in its extent throughout the genome, with the proportion of methylated CpGs much lower in invertebrates: 0-14% as compared to 60-90% in mammals (Bewick *et al.* 2017, Glastad *et al.* 2011). The evolutionary reasons for these differences are not yet clear, although it is thought that gene body methylation is an ancient feature of eukaryote genomes (Zemach *et al.* 2010). However, DNA methylation status influences how genes (and phenotypes) are expressed in many organisms, including reproductive vs non-reproductive worker bumblebees (Amarasinghe *et al.* 2014), caste differentiation in honey bees (Kucharski *et al.* 2008, Lyko *et al.* 2010), transgenerational response to the presence of predators in *Daphnia magna* (Schield *et al.* 2016), and between lateral plate morphotypes in Threespine Stickleback (Smith *et al.* 2014). Therefore, DNA methylation patterns are very much part of mapping genotypes to phenotypes.

The flexible control of gene expression provided by mechanisms such as DNA methylation has also caught the interest of biologists interested in the underlying mechanisms of phenotypic plasticity, and how such plasticity may evolve (Bonduriansky and Day 2009, 2018). Moreover, there has also been great interest in the extent to which epigenetic modifications may be inherited across generations (Bonduriansky and Day 2009, Burggren 2016, Lind and Spagopoulou 2018). Transmission of epigenetic state allows transmission of parent-of-origin information, or so-called genomic imprinting, setting the scene for genomic conflict (Haig 2002, Patten *et al.* 2014). Genomic conflict arises when maternally- and paternally-inherited alleles have different evolutionary optima for traits expressed in an individual (Haig 1997, 2002). A classic example is the expression of maternally- or paternally-inherited alleles during embryonic development in mammals: paternally-inherited alleles are selected to extract more resources from the mother *in utero* compared to maternally-inherited alleles (Moore and Haig 1991). This conflict is now known to be, at least in part, facilitated by parent-of-origin specific gene expression of insulin-like growth factor II (Igf2) and Igf2r. In mammals, the Igf2 gene promotes growth and cellular differentiation during development, and also regulates the placental supply of nutrients and the demand of nutrients by the foetus (Constancia *et al.* 2002).

To further understand the role epigenetic modifications such as DNA methylation have in influencing phenotypes and their evolution, we need robust ways of manipulating them. Here we consider the efficacy of the chemical 5-aza-2’-deoxycytidine (5-aza-dC) as a way of manipulating DNA methylation, using the parasitoid wasp *Nasonia vitripennis*. The extent and nature of DNA methylation across the *N. vitripennis* genome has recently been characterised (Wang *et al.* 2013, Pegoraro *et al.* 2016), and the presence in the genome of a full “methylation toolkit” has also been confirmed (Werren *et al.* 2010). 5-aza-dC is well-known from the cancer literature as a hypomethylating agent and is used in the treatment of acute myeloid leukaemia and myelodysplastic syndrome (Bryan *et al.* 2011, Momparler 2012, Seelan *et al.* 2018). In cancer patients, many genes that suppress leukemogenesis in humans are silenced by aberrant DNA methylation. 5-aza-dC acts as a demethylating agent to reactivate these genes (see below for details of action). In recent years, 5-aza-dC has been used across a range of species as an experimental tool to explore how DNA methylation influences phenotypes including diapause in *Nasonia vitripennis* (Pegoraro *et al.* 2016) and worker reproductive status in bumble bees (Amarasinghe *et al.* 2014). Previously, we used 5-aza-dC to explore whether DNA methylation was involved in the control of facultative sex allocation in *N. vitripennis* (Cook *et al.* 2015).

Like all Hymenoptera, *N. vitripennis* is haplodiploid, with males arising from unfertilised haploid eggs and females arising from fertilised diploid eggs. Females have putative control over the sex of their offspring, releasing sperm to produce daughters or not to produce sons. Female *Nasonia* allocate sex broadly in line with the predictions of Local Mate Competition theory (Hamilton 1967, Taylor and Bulmer 1980, Werren 1980, Werren 1983, Shuker and West 2004, Burton-Chellew *et al.* 2008; reviewed by West 2009). Females laying eggs alone on their blowfly pupae hosts produce very female-biased offspring sex ratios to minimise competition amongst her sons for mating opportunities. As more females (foundresses) lay eggs on the same host(s) together, females produce less female-biased sex ratios, as LMC is reduced and the fitness gained through the production of sons and daughters moves towards equality (Hamilton 1967). When treated with 5-aza-dC, the facultative sex allocation response in female *Nasonia* was maintained, suggesting that this plastic behaviour in and of itself did not require certain patterns of methylation. Rather, the sex ratios (as proportion male) produced by females were shifted slightly upwards, with more sons produced by treated females than by controls. This subtle shift is as predicted by genomic conflict over sex allocation theory, which requires some form of genomic imprinting (Wild and West 2008). Assuming that DNA methylation is the mechanism by which parent-of-origin allelic information is carried in *Nasonia* (although see Wang *et al.* 2016), our data therefore provided the first (albeit indirect) evidence for genomic conflict influencing sex allocation (Cook *et al.* 2015a).

Recently, Ellers *et al.* (2019) challenged the effects of 5-aza-dC as a general modifier of methylation status, considering CpGs in nine genes in *Nasonia vitripennis*. Ellers *et al.* (2009) failed to find changes in methylation status across any of the 155 CpGs in their study, questioning the action of 5-aza-dC in *Nasonia*, and hence of course the interpretation of the effects of 5-aza-dC in our earlier study (Cook *et al.* 2015a). Fully negative results are of course very hard to interpret, but Ellers *et al.* (2009) nonetheless raised valid concerns of the efficacy of the chemical. As noted above, levels of methylation in insects are generally low, and a genome-wide study would provide greater resolution to explore the effects of this chemical in an ecologically valuable model system. Here we consider the effects of 5-aza-dC across the whole genome using a Bisulfite sequencing (BS-Seq) experiment. There is evidence to suggest that the effects of 5-aza-dC are unlikely to be random across methylated CpGs in a given genome. For instance, in mammals, genomic context can influence the extent to which 5-aza-dC is effective as a demethylating agent. DNA sequence context, the distribution of transcription factor binding sites, and chromatin structure are known to influence the DNA methylation patterns produced by 5-aza-dC exposure (Mossman *et al.* 2010, Hagemann *et al.* 2011, Ramos *et al.* 2015). Although patterns of methylation differ among eukaryotes, the epigenetic modification itself is chemically identical. Here, we verify that 5-aza-dC influences patterns of methylation across the genome in *N. vitripennis*. Altered methylation patterns produced by 5-aza-dC exposure were non-random, with enrichment in genes associated with transcription-factor activity and sequence specific binding.

## Results

Treatment with 5-aza-dC had wide-ranging effects on DNA methylation across the genome. First, samples clustered strongly by 5-aza-dC exposure regime at the gene-level in both t-SNE visualisation (Fig 1) and in principle component analyses (Fig 2), indicating that 5-aza-dC has a strong effect on the methylation of genes. Both analyses show strong clustering, and that wasps exposed to 5-aza-dC for 24 hours are more like controls than wasps exposed to 5-aza-dC for 48 hours. In addition, the PCA analysis showed clear separation along PCA1 (which presumably is heavily associated with exposure regime; Fig 2). Importantly though, this analysis also emphasises the extent of variation in response across the three treatment regimes, which is smallest for the controls, and largest for the 48 hour-exposed wasps (note the spread along PCA axis 2 in Fig 2). In addition, both cluster analyses placed the controls as some way intermediate between the two 5-aza-dC exposure treatments, which we consider further in the next two paragraphs. Both analyses also show that exposure regime dominated the clustering, over and above of any effects of either tissue-type (wasp heads vs bodies) or collection time (hours post-exposure), suggesting that this is the key effect. When we consider clustering at the level of individual CpG loci, clustering is much less obvious, using both t-SNE and PCA. However, the control treatment appears to cluster within the others in the t-SNE analyses, and the 24-hour and 48-hour treatments do separate to some extent, e.g. with the 24h samples more to the top and right of Fig S1 and the 48h samples more to the bottom and left (see Figs S1 and S2). The differences between the individual CpG and gene-level cluster analyses are likely to be due to a combination of low coverage and a greater stochasticity of methylation at the individual CpG level (see discussion).

**Fig 1.**
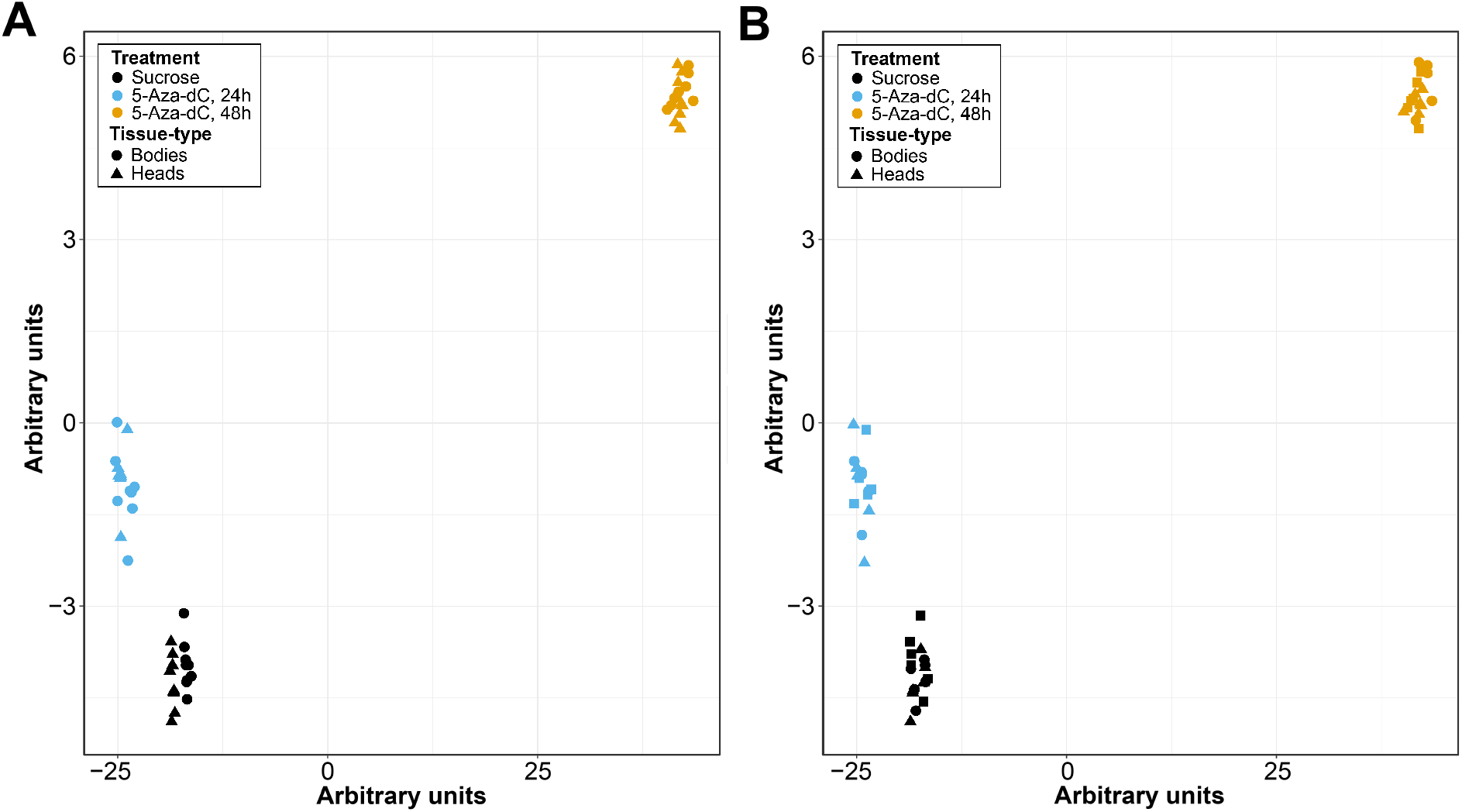
t-SNE of the proportion of methylated reads per gene. Shows the effects of (A) Exposure regime and tissue type and (B) Exposure regime and harvest time.

**Fig 2.**
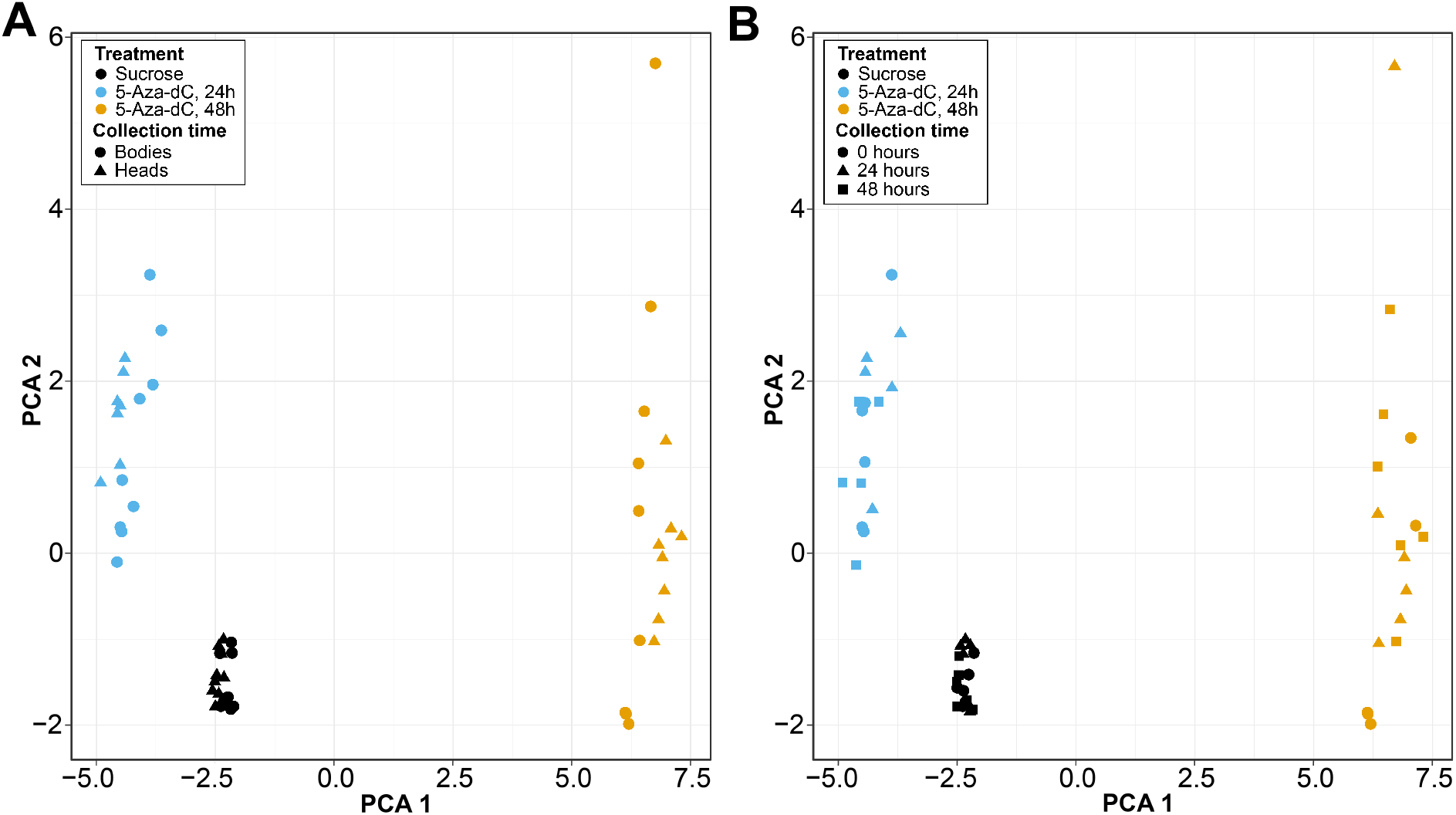
PCA of the proportion of methylated reads per gene. Shows the effects of (A) Exposure regime and tissue type and (B) Exposure regime and harvest time.

Second, a more in-depth examination of the data using a generalised linear modelling (GLM) approach to examine differential methylation at the gene level found that 5-aza-dC exposure had a significant effect on the methylation status of the majority of genes examined; exposure treatment was significant as a main effect at an FDR level of 5% for 8,725 out of 11,582 genes, or 75.3% of genes (see also Fig S3). Moreover, both tissue-type and collection time also influenced methylation status across a large number of genes (significant as main effects in the GLMs for 3833 and 3566 genes respectively). Significant interactions between factors were also present for the majority of genes (exposure regime * collection time = 10,776 genes, exposure regime * tissue type = 10,484 genes, collection time * tissue type = 5,364 genes, and exposure * collection time * tissue type = 10,484 genes), showing widespread evidence for significant interplay between these factors. These results emphasise the context-dependent nature of the action of 5-aza-dC. Full results from the GLMs are presented in Table S1.

Interestingly, it appears that methylation tends to increase initially in response to exposure to 5-aza-dC, an effect that is noticeable after 24h of exposure, before decreasing after 48h of exposure (Fig 3A). This means that the length of time individuals are exposed to the chemical may well influence what kind of methylation changes are observed; in other words, both hyper- and hypo-methylation can result from 5-aza-dC treatment. In terms of when wasps were collected after exposure, 3,566 genes displayed significantly altered methylation in association with collection time, with a more pronounced decrease in methylation in samples collected 48h post-exposure relative to 24h post-exposure (when compared to 0h post-exposure; Fig 3B). This suggests that the effect of 5-aza-dC persists at least 48h after exposure has stopped. Finally, 3,833 genes displayed significantly altered methylation patterns in association with tissue type (Fig 3C), with lower methylation overall in the head. However, we re-iterate that these main effects need to be contextualised within the pattern of widespread interactions seen in the data, with 91% of genes exhibiting a significant second-order interaction between exposure, tissue type and collection time.

**Fig 3.**
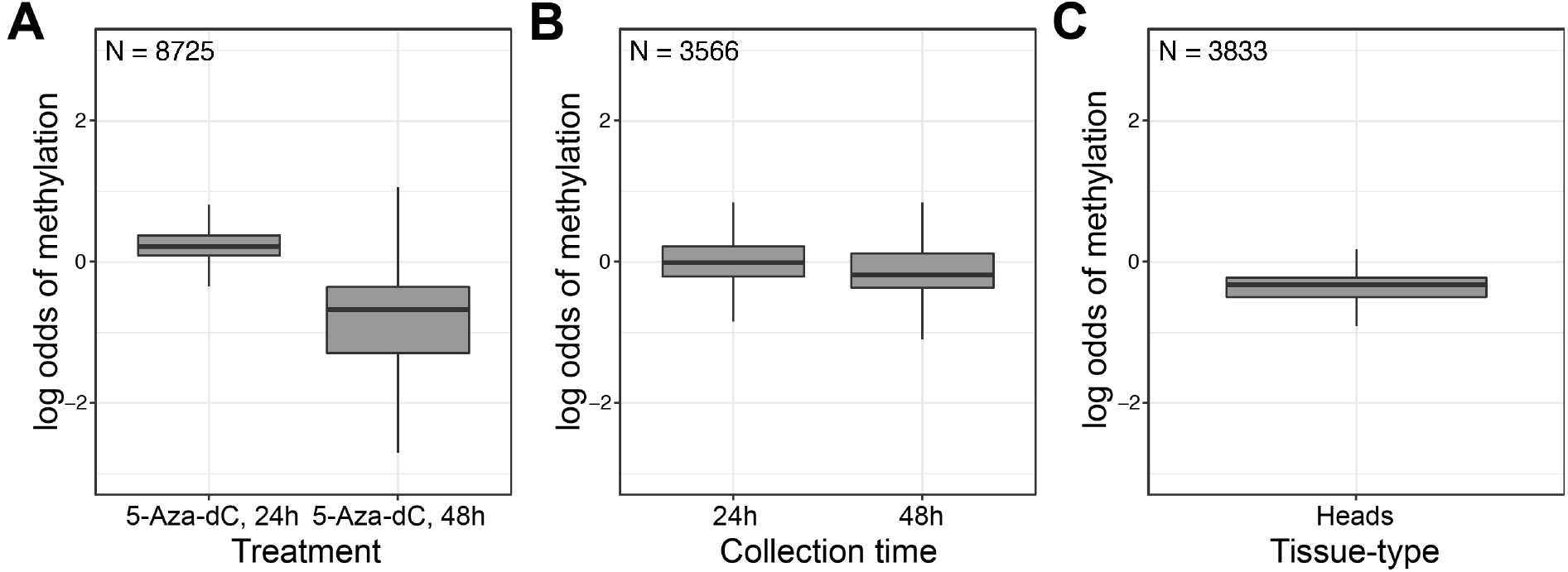
Gene methylation status displayed as log odds of methylation for each main effect in the experiment. A significant effect on gene methylation status was noted in response to (A) 5-aza-dC exposure regime (relative to sucrose control), (B) harvest time after exposure (relative to zero hours) and (C) tissue-type (heads relative to bodies). N= number of genes with significantly differentially methylated CpGs in association with each main effect.

Gene-set enrichment analysis, using *N. vitripennis* GO terms, when genes were ranked by their response to exposure regime is presented in Table S2. Broadly, we identified 169 significantly enriched GO terms (p < 0.05), and 80 of these terms fell under the “Biological process” (BP) category, 18 under “Cellular component” (CC), and 71 under “Molecular function” (MF). The top five enriched GO terms were “sequence-specific DNA binding” (GO:0043565), “regulation of transcription, DNA templated” (GO:0006355), “ATP binding” (GO:0005524), “DNA binding transcription factor activity” (GO: 0003700), and “protein kinase activity” (GO:0004672) (Table S2). When the analysis was repeated using GO terms for the *D. melanogaster* orthologs, a higher number of significantly enriched GO terms was obtained (700; 538 BP, CC 71, MF 91; Table S3). The top five GO terms for the *D. melanogaster* analysis were “nucleus” (GO:0005634), “positive regulation of transcription by RNA polymerase II” (GO:0045944), “regulation of Notch signalling pathway” (GO:0008593), “imaginal disc-derived wing morphogenesis” (GO:0007476), and, making a reappearance, “sequence-specific DNA binding” (GO:0043565). This analysis combined GO-terms when *N. vitripennis* genes had multiple *D. melanogaster* orthologs, which could lead to bias. H, however, we retrieve very similar enriched terms when we used only the GO terms that were shared between the multiple orthologs (Table S4), suggesting any bias caused by this approach is small.

## Discussion

Our genome-wide analysis of CpGs across more than 11,000 genes showed that the demethylating agent 5-aza-dC has very widespread effects on patterns of DNA methylation in the *Nasonia vitripennis* genome. The methylome has been characterised in *Nasonia vitripennis* by Wang *et al.* (2013) and Beeler *et al.* (2014), and in common with other insect species, DNA methylation appears to be primarily associated with gene bodies (including intron-exon boundaries), and a rather low proportion of CpGs are methylated. As mentioned above, previous work that has attempted to confirm the effects of 5-aza-dC on DNA methylation have had mixed results. Pegoraro *et al.* (2016) confirmed the effects of 5-aza-dC across five genes (four via a qPCR MethylQuant assay and one via cloning a fragment of the *msn* gene from treated versus untreated individuals), whilst Ellers *et al.* (2019) have recently failed to show any effects of 5-aza-dC on the methylation status of 155 CpGs across nine genes (they used high-throughput bisulfite amplicon sequencing (BSAS); see Redshaw *et al.* (2014) for a discussion of different methylation quantification assays). Our larger-scale study has both confirmed the action of 5-aza-dC as a disruptor of methylation in *Nasonia*, but also suggested why it might be so difficult to pin down those effects. In particular, we find that whilst we see statistically significant main effects of exposure regime on DNA methylation (which includes differences in length of exposure), we also see effects of tissue type (albeit at the crude level of heads versus bodies), and time since exposure to 5-aza-dC. Moreover, we also see many statistically significant interactions between these factors; all told, the action of 5-aza-dC is likely to be context-dependent, both in terms of the sequences more likely to be targeted but also in terms of the experimental details, including target tissues. Nonetheless, our results strongly suggest that 5-aza-dC is not chemically inert in terms of influencing DNA methylation in *Nasonia* – indeed, quite the opposite.

That the effects of 5-aza-dC are non-random in terms of the genes it influences are further confirmed by our gene ontology enrichment analyses. We see enrichment for genes associated with DNA binding, regulation of transcription, and transcription factor activity. These processes are of interest given that we might expect the regulation of transcription to be a crucial component of the epigenetic role of DNA methylation, especially given what we know from both mammals and insects, even though the targets (throughout the genome versus exons) and outcomes (transcriptional silencing versus stable gene expression) are typically different in the two taxa. Our analysis certainly confirms the sequence-specific action of 5-aza-dC seen in other species.

A further important effect we see is that 5-aza-dC exposure can result in hyper-methylation as well as hypo-methylation. This means that the characterisation of 5-aza-dC as a strictly demethylating agent may be somewhat misleading. The complexity of outcomes of 5-aza-dC in mammals, in particular in terms of trying to understand its action for cancer treatments, is becoming increasingly well-appreciated (Seelan *et al.* 2018). As such, our data confirm in an insect what is being seen in mammalian cells. We are not well placed to speculate on the mechanism(s) by which 5-aza-dC may lead to hyper-methylation (see instead Seelan *et al.* 2018), even though its hypomethylating effect is well characterised chemically (see Introduction). However, if 5-aza-dC disrupts the regulation of the DNA methylation machinery itself, for instance in terms of how the Dnmt genes are expressed and act, then the downstream outcomes could be both hyper- and hypo-methylation. Perhaps related to this, in their study of DNA methylation and the control of the diapause response in *Nasonia*, Pegoraro *et al.* (2016) used both 5-aza-dC and RNA interference (RNAi) to manipulate methylation, in the latter case disrupting DNA methylation by knocking down both Dnmt1a and Dnmt3. Whilst both exposure to 5-aza-dC and the RNAi constructs affected how diapause was influenced by exposure to short or long day-length, they did so in different ways. Whilst both manipulations abolished the day-length response, the RNAi of Dnmt1a and Dnmt3 resulted in a change in how wasps responded to long-day length (increasing the diapause response), whilst 5-aza-dC led to a change in response to short-day length (decreasing the diapause response). As such, different manipulations of DNA methylation may lead to different phenotypic outcomes, depending on the regulatory causes and consequences of the methylation pattern itself. More generally then, it is clear that much remains to be elucidated in how 5-aza-dC shapes methylation patterns, in both insects and mammals.

On a more technical note, perhaps unsurprisingly we found more significant genes in the GO enrichment analysis using the GO terms for *Drosophila melanogaster* orthologs. This is perhaps because of the more detailed gene ontology information available for the *D. melanogaster* genome. Nonetheless, the two analyses provide a similar story in terms of GO enrichment. In addition, the majority of our analyses were undertaken at the level of the gene. Perhaps given the variation in coverage at the level of individual CpGs, the clustering analyses were not by any means as clear cut as the gene-level results. However, in the t-SNE analysis (Fig S1), the control replicates cluster together, albeit within the broader distribution of the other two exposure treatment groups. The effect of genomic and sequence context may be particularly relevant here: our analysis at the level of the gene (see Methods for details) brings together CpGs that share a lot of context, by virtue of being associated with a given genetic locus. The differences in the clustering at the individual CpG versus gene levels may therefore represent the fact that the action of 5-aza-dC does appear to be non-random, and that we gain clarity when we consider closely associated CpGs whose patterns of methylation, and responses to 5-aza-dC, are more similar than randomly chosen CpGs across the genome.

In conclusion, our results suggest that 5-aza-dC remains a potential tool for evolutionary ecologists wishing to explore how DNA methylation influences phenotypic expression. As with all genomic approaches however, no one approach gives all the answers. 5-aza-dC may prove a very useful, if rather blunt, tool for first assessing the range of phenotypes that are influenced by methylation status, whilst no doubt more targeted knock-downs and genetic manipulations will be needed to link phenotypes to causal CpGs.

## Materials and Methods

### Study species

*Nasonia vitripennis* (Hymenoptera, Chalcicoidea) is a generalist parasitoid of large dipteran pupae including species of Calliphoridae. Females oviposit between 20 and 50 eggs in an individual host, with male offspring emerging just before females (after approximately 14 d at 25 °C: Whiting 1967). Males are brachypterous and unable to fly, remaining close to the emergence site where they compete with each other for emerging females, including their sisters. Females disperse after mating to locate new hosts. The females used in this experiment were from the wild-type AsymC strain, originally isolated in 1986 by curing the wild-type strain LabII of *Wolbachia*. Wasps have been maintained on *Calliphora vomitoria* or *C. vicina* hosts at 25 °C, 16L:8D light conditions ever since, and AsymC is also the reference genome strain (Werren *et al*. 2010). The quantitative genetic basis of sex ratio variation has been repeatedly quantified (Pannebakker *et al.* 2008, 2011), whilst whole-genome transcription studies have confirmed that facultative sex allocation takes place without discernible changes in gene expression (Cook *et al.* 2015b, 2018).

### 5-aza-2’-deoxycytidine

We used 5-aza-dC (Sigma Aldrich Company Ltd., Gillingham, UK) at a non-lethal concentration (10 μM). Widely used in epigenetic anticancer treatments (Seelan *et al.* 2018), 5-aza-dC is a nucleoside analogue that works by inhibiting DNA methyltransferase activity (Piskala and Sorm 1964, Jones and Taylor 1980, Momparler 1985, Christman 2002). When initially introduced to a cell, 5-aza-dC is essentially inert but is then converted by processes inherent in the cell to its active form 5-aza-dCTP (Seelan *et al.* 2018). This active form is readily incorporated into DNA during S-phase of the cell cycle in the place of cytosine. In the daughter strands arising from a replication event, DNA methyltransferase 1 (Dnmt1) will attempt to restore the patterns of methylation that were present in the parent strand. However, Dnmt1 becomes irreversibly bound to the 5-aza-dCTP residues in the daughter strands, resulting in a diminishing pool of Dnmt1. Thus, administering 5-aza-dC results in a passive loss of DNA methylation, the key effect of 5-aza-dC (for a fuller description, see Cook *et al.* 2019).

### Experimental design

To control for possible host and other maternal effects, experimental females were not drawn straight from stock populations. Instead, two-day-old, mated, wild-type AsymC females were isolated from the mass cultures into individual glass vials. Each female was provided with three hosts and allowed to oviposit. Experimental females were drawn from the F1 generation, two days after emergence, one female per “grandmother”. We then employed a three by three factorial design whereby females were allocated to one of three 5-aza-dC exposure regimes and were harvested post-feeding at three time points. Exposure regimes were a) 20% sucrose for 24h, b) 20% sucrose supplemented with 10 μM 5-aza-dC for 24h and c) 20% sucrose supplemented with 10 μM 5-aza-dC for 48h. For each of the three regimes, females were harvested at 0, 24 or 48h post-exposure. Throughout the experiment, females were kept individually in glass vials and incubated at 25 °C, 16L:8D. Solutions (200 μl) were provided in the lid of a 2 ml microcentrifuge tube. For individuals exposed to 5-aza-dC for 48 h, the solutions were topped up by 100μl half-way through the exposure period to compensate for evaporation. All solutions were stained green with food colouring such that a subset of 10 females from each treatment combination could be dissected to check that feeding had occurred. At the point of harvest, females were flash-frozen in liquid nitrogen and stored at −80 °C prior to DNA extraction.

### DNA extraction and bisulphite-conversion

Prior to DNA extraction, individuals were removed from −80 °C storage and heads were quickly excised from bodies using a sterile scalpel and a cryloyser to keep samples cold. Heads (and likewise bodies) from the same treatment combination were randomly pooled into groups of 10 to give a maximum of 3 biological replicates for each treatment combination giving a total of 54 DNA samples. Briefly, tissue was homogenised using a micropestle in 350 μl CTAB (Hexadecyltrimethylammonium Bromide) buffer with subsequent overnight proteinase K digestion (400 μg) at 56 °C. Samples were cooled to room temperature prior to RNase A digestion for 1 hour at 37 °C. A chloroform: isoamyl-alcohol wash (300 μl) was performed twice before ethanol/sodium acetate precipitation of the DNA. DNA purity was analysed using the Nanodrop spectrophotometer with all samples having a 260/280 ratio of ≥1.8. DNA integrity was confirmed using agarose gel electrophoresis.

### Bisulphite sequencing

Bisulphite conversion, library preparation and sequencing were carried out by Edinburgh Genomics. Briefly, bisulphite conversion was carried out using the EZ DNA Methylation Kit (Zymo Research, Irvine USA), designed to reduce template degradation during the harsh bisulphite treatment whilst ensuring complete conversion of the DNA. Library preparation was carried out using the TruSeq DNA methylation Kit which takes the single-stranded DNA that results from bisulphite treatment and converts it into an Illumina sequencing library. Sequencing of bisulphite-treated DNA was carried out on the Illumina HiSeq 2500 in high output mode with 125bp paired-end reads. Library preparation failed for three of the samples due to low levels of input DNA, leaving only two biological replicates for three out of 18 treatment combinations and a total of 51 Bisulphite-sequenced samples (Table 1).

**Table 1.** A summary of the experimental design^a^.

| Treatment Combination | 5-aza-dC exposure regime | Harvest time | Tissue type | No. replicates (1 x rep. = 1 pool of 10) |
| --- | --- | --- | --- | --- |
| 1 | 20% sucrose for 24 h | 0h | Bodies | 3 |
| 2 | 20% sucrose for 24 h | 0h | Heads | 3 |
| 3 | 20% sucrose for 24 h | 24h | Bodies | 3 |
| 4 | 20% sucrose for 24 h | 24h | Heads | 3 |
| 5 | 20% sucrose for 24 h | 48h | Bodies | 3 |
| 6 | 20% sucrose for 24 h | 48h | Heads | 3 |
| 7 | 10µM 5-aza-dC for 24 h | 0h | Bodies | 3 |
| 8 | 10µM 5-aza-dC for 24 h | 0h | Heads | 3 |
| 9 | 10µM 5-aza-dC for 24 h | 24h | Bodies | 3 |
| 10 | 10µM 5-aza-dC for 24 h | 24h | Heads | 2 |
| 11 | 10µM 5-aza-dC for 24 h | 48h | Bodies | 3 |
| 12 | 10µM 5-aza-dC for 24 h | 48h | Heads | 2 |
| 13 | 10µM 5-aza-dC for 48 h | 0h | Bodies | 3 |
| 14 | 10µM 5-aza-dC for 48 h | 0h | Heads | 2 |
| 15 | 10µM 5-aza-dC for 48 h | 24h | Bodies | 3 |
| 16 | 10µM 5-aza-dC for 48 h | 24h | Heads | 3 |
| 17 | 10µM 5-aza-dC for 48 h | 48h | Bodies | 3 |
| 18 | 10µM 5-aza-dC for 48 h | 48h | Heads | 3 |

<sup>a</sup> Female wasps were exposed to 1 of 3 5-aza-dC exposure regimes including a control. Females were then harvested at 1 of 3 timepoints after the exposure period heads were excised from bodies to give a degree of tissue-specificity to the analysis. Three biological replicates were available for almost all treatment combinations with each biological replicate consisting of tissue from three *N. vitripennis* females.

### Raw data processing

Initial data processing was carried out by Edinburgh Genomics. Briefly, reads were trimmed using Cutadapt version 1.12 removing any adapter sequences and for quality using a threshold of 30. After trimming reads were required to have a minimum length of 50 bp. *Nasonia vitripennis* genome version 2.1 and the associated annotation, available for download at Ensembl (http://metazoa.ensembl.org/Nasonia_vitripennis/Info/Index), were used as a reference. Reads were aligned to the reference genome using Bismark (version 0.16) with the parameter “bowtie2”. PCR duplicates were removed from the resulting BAM files using samtools with the parameter “view -F 1024”. Data for each CpG was then extracted from the BAM files using the Bioconductor package methylKit (version 0.99.3) with R (version 3.3.1; R Core Team 2017). The methylation status of each CpG for all samples was calculated using the function “processBismarkAln” using the default parameters, except “min.cov=1 no;ap=TRUE”, which excludes CpG’s not covered by any reads across all samples and does not count CpGs covered by overlapping ends of a paired read as covered twice. The 1% of bases with the highest coverage were removed using the function “filterByCoverage” to eliminate potential PCR bias. After initial processing as outlined above, the raw data comprised percentage methylation at 4,150,376 CpG loci (no. of reads in which the CpG is methylated/total no. of reads covering the CpG) for each of 51 samples. CpG read counts were then summed by gene, (including 1Kb up and downstream) using gene annotation from Ensembl (version 2.1. available from: ftp://ftp.ensemblgenomes.org/pub/metazoa/release-40/gff3/nasonia_vitripennis). Around half (2,119,643) of the CpG’s in the full dataset fell within genic regions with 14,765/17,279 genes (85%) having at least one CpG present. Median coverage across the whole experiment (N = 51 BS-seq libraries) was 898X, but coverage was heterogeneous between loci (Fig 4) and samples (Table S5). Mean read coverage per sample ranged from 2.6X to 48.2X. Four samples had less than 10X mean coverage per CpG, however, these samples were distributed across treatment groups.

**Fig 4.**
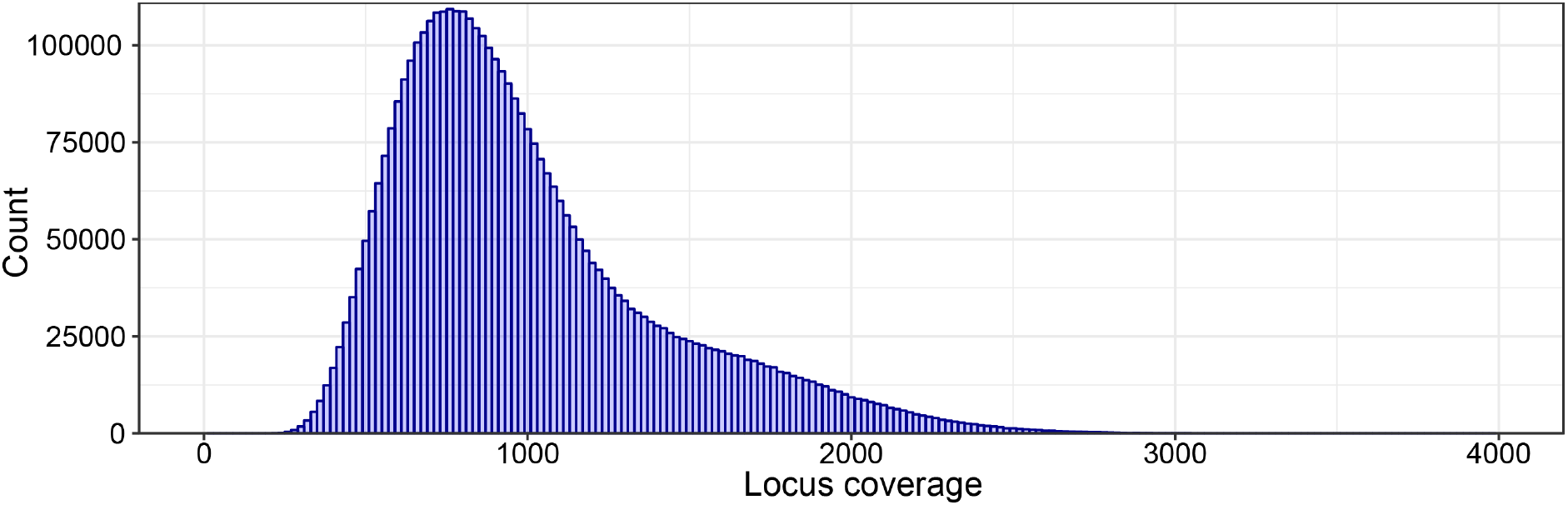
Read coverage by locus across all 51 BS-Seq libraries.

### Statistical analysis

The overall structure of our dataset in terms of the proportion of methylated reads per gene and per loci was visualised with two complementary analyses: a principle component analysis (PCA), which allows samples to be visualised in relation to the two major axes of variation in our experiment, and a T-distributed Stochastic Neighbour Embedding (t-SNE) analysis. t-SNE is a machine-learning algorithm for data visualisation well-suited for high-dimensional datasets (van der Maaten and Hinton 2008). The PCA was conducted using the R base function prcomp with default values. To perform the t-SNE analysis we used the R package Rtsne (Krijthe 2015), with default values and perplexity set to 15. For the PCA and t-SNE analyses any genes or loci with 0 coverage in any one sample were discarded prior to analysis.

To examine the effect of 5-aza-dC exposure on individual genes, we modelled the number of methylated and unmethylated reads from a gene as a generalized linear model (GLM) with a binomial distribution with the following terms: exposure, tissue-type, collection time and their interactions. Genes with 0 coverage for all samples in any particular treatment group were excluded from our analysis (leaving a total of 11582 genes). Significance of each term was determined by a likelihood ratio test, which was corrected for multiple testing using Benjamini and Hochberg’s algorithm (Benjamini and Hochberg 1995), with statistical significance set to 5%. Coefficients (log odds of methylation) for each term were extracted from the full model. All statistical analyses were conducted in R (v. 3.4.1; R Core Team 2017).

### Functional enrichment

*N. vitripennis* GO-terms were downloaded from “hymenopteramine” (http://hymenopteragenome.org/hymenopteramine/begin.do) on 27^th^ September 2018. In addition, we used hymenopteramine to firstly obtain *D. melanogaster* orthologs for each *N. vitripennis* gene, from which we could then obtain the associated *D. melanogaster* GO terms. We used both of these GO-term sets for enrichment analyses (below). Some *N. vitripennis* genes had multiple *D. melanogaster* orthologs. In these cases, all GO-terms from each *D. melanogaster* ortholog were combined together. To ensure this did not bias our results we repeated our analyses, but instead of combining GO-terms we kept only GO-terms that were shared between all orthologs.

To examine which processes are most affected by 5-aza-dC exposure, we conducted gene set enrichment analyses using the R package TopGO (v. 2.28.0; Rahnenfuhrer 2016) using the elim algorithm to account for the GO topology. Gene set enrichment analyses identify enriched GO terms in a threshold-free way, by finding GO-terms that are overrepresented at the top of a ranked list of genes. Here we ranked genes by FDR for the exposure effect. GO terms were considered to be significantly enriched when p < 0.05.

## Acknowledgments

This work was initiated as part of NERC grant NE/J024481/1. We are very grateful to Mark Lammers and Jacintha Ellers for their collegiate and constructive engagement with our previous DNA methylation work. We are also grateful to Matthew Arno and the Edinburgh Genomics team for their sequencing and downstream support. Raw data are available from the Gene Expression Omnibus hosted by the National Center for Biotechnology Information (Accession: GSE125388). Scripts for the analyses in this paper are available at: https://github.com/DarrenJParker/5-Aza-dC_methylation.

## Notes

#### Summary of Updates

Updated ms. Fixed figure legend.

