## Supplementary material for "Genome-wide disruption of DNA methylation by 5-aza-2’-deoxycytidine in a parasitoid wasp": Supp Figs

### Supporting Information

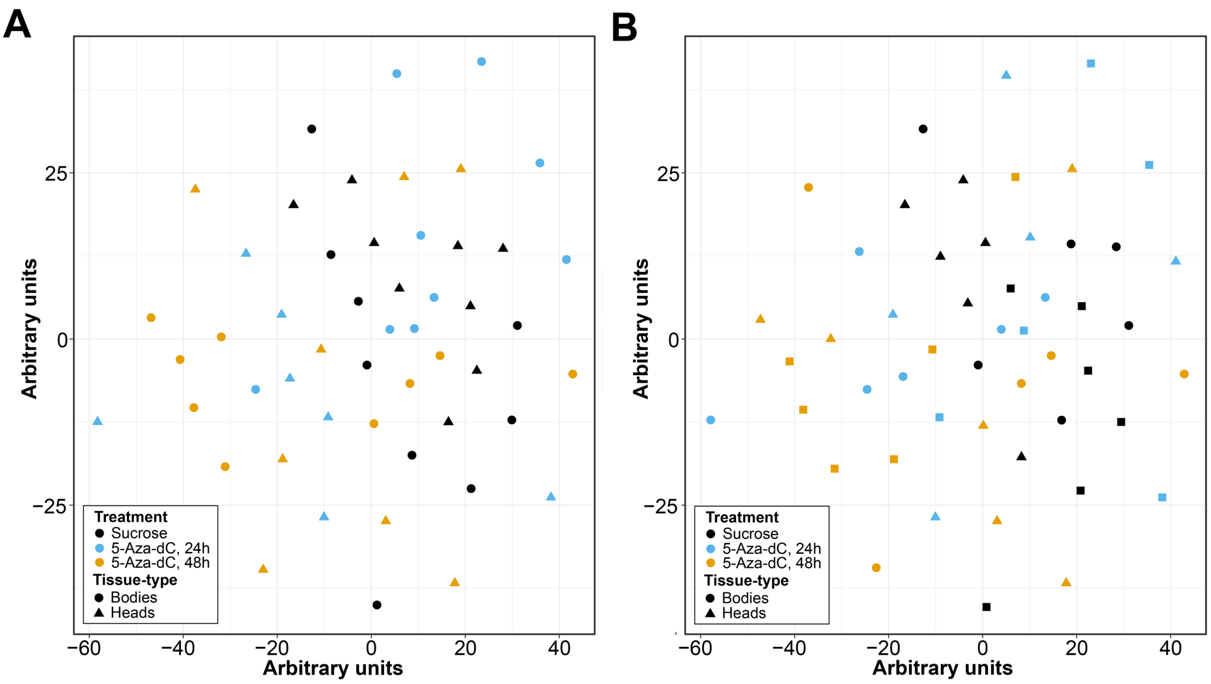


**Fig S1. t-SNE of the proportion of methylated reads per CpG locus**. Shows the effects of (A) Exposure regime and tissue type and (B) Exposure regime and harvest time.

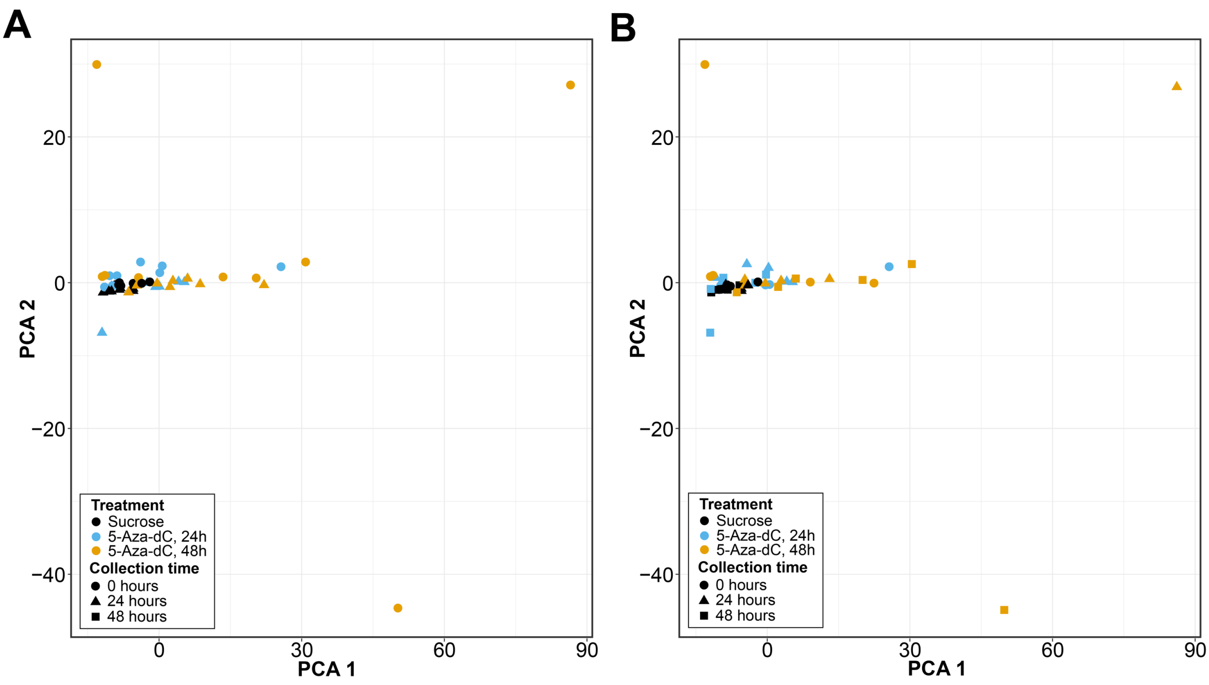


**Fig S2. PCA of the proportion of methylated reads per CpG locus.** Shows the effects of (A) Exposure regime and tissue type and (B) Exposure regime and harvest time.

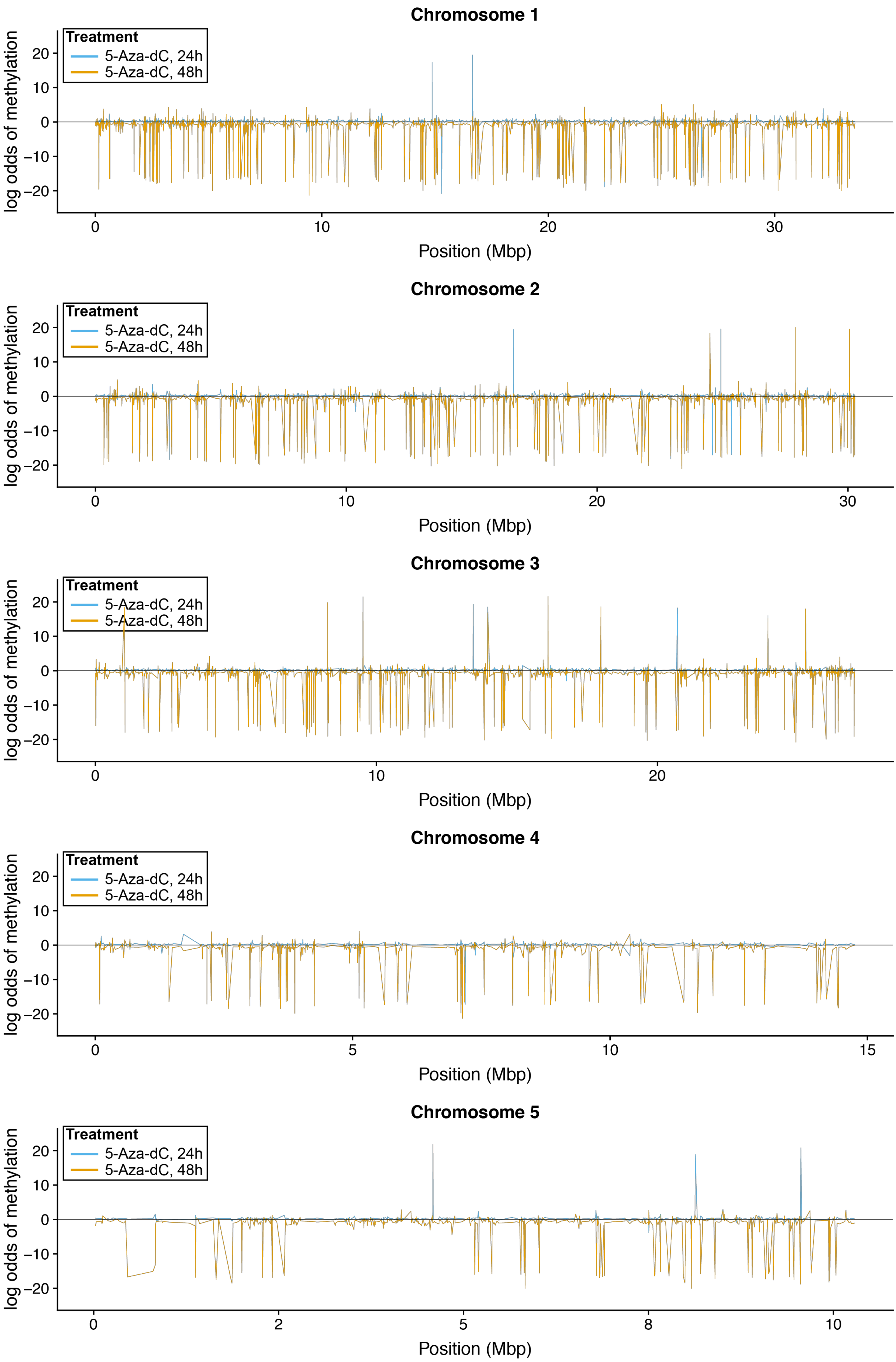


**Fig S3. Effect of 5-aza-dC exposure regime on gene methylation status (log odds of methylation) for each gene ordered along each chromosome.**

**Table S1. Number of reads and coverage of each sample.**

**Table S2. Output of coefficients and significance values from the GLM analysis.**

**Table S3. Enriched GO terms for treatment using *Nasonia vitripennis* GO terms.**

**Table S4. Enriched GO terms for treatment using *D. melanogaster* GO terms, when GO-terms from multiple orthologs are combined together.**

**Table S5. Enriched GO terms for treatment using *D. melanogaster* GO terms, when only GO-terms which are the same are kept (when a *Nasonia vitripennis* gene has multiple *D. melanogaster* orthologs).**
